# Ecosystem engineers cause biodiversity spill-over: beavers affect breeding bird assemblages on both wetlands and adjacent terrestrial habitats

**DOI:** 10.1101/2024.04.24.591038

**Authors:** Izabela Fedyń, Wojciech Sobociński, Sławomir Czyżowicz, Jakub Wyka, Michał Ciach

## Abstract

The influence of ecosystem engineers on habitats and communities is commonly acknowledged in a site-bounded context, i.e. in places directly affected by the presence of the focal species. However, the spatial extent of the effects of such engineering is poorly understood, raising the question as to what impact they have on ecosystems situated beyond the species’ direct influence. Beavers *Castor* spp., iconic ecosystem engineers, are capable of significantly transforming aquatic ecosystems. Their presence boosts biodiversity in near-water habitats, but as a result of cascading processes, beavers may affect terrestrial habitats situated beyond the range of their immediate activity. Our study investigates the breeding bird assemblage along a spatial gradient from the water to the forest interior on central European watercourses modified and unmodified by beavers. The results show that beaver sites are characterized by a higher species richness and abundance of breeding birds than unmodified watercourses. Such sites also host a different species pool, as 27% of the recorded bird species occurred exclusively on the beaver sites. The effect of the beaver’s presence on the bird assemblage caused this to spill over from the wetlands to adjacent terrestrial habitats located up to 100 m from the water’s edge, where the species richness and abundance was higher and the species composition was substantially modified. We also found a positive correlation between the total area of beaver wetland and the numbers of bird species and individuals recorded. Our study indicates that beaver activity has major consequences for ecosystem functioning that extend beyond the wetland, spilling over on to adjacent terrestrial ecosystems. These findings add to the general understanding of the spatial context of the ecosystem engineering concept, as the changes brought about by engineers have an influence beyond the area of their immediate occurrence. Our work also has implications for landscape planning and management, where existing beaver sites with terrestrial buffer zones may constitute a network of biodiversity hot-spots.

**Highlights:** • Beavers create hot-spots of bird richness and abundance in temperate forests

• The beaver’s impact spills over beyond wetlands on to terrestrial ecosystems

• Bird richness, number and composition is changed up to 100 m from beaver sites

• The area of a beaver wetland positively correlates with bird richness and numbers

• The beaver is an umbrella species for the riparian forest bird community

## Introduction

Ecosystem engineers, with their remarkable ability to modify the environment, have a significant impact in shaping the diversity and composition of communities within the area of their occurrence (Jones et al., 1994). Globally recognized as a facilitative process, ecosystem engineering contributes to a 25% increase in biodiversity (Romero et al., 2015). Ecosystem engineers not only reshape the structure and resources of the environment, but their activities also have ecosystem-level consequences by modulating interactions within communities, as habitat modification often leads to changes in species composition (Losapio et al., 2023). The concept of involving species with specialized abilities to modify the environment is shifting more and more often from the theoretical framework to practical conservation applications, such as restoration projects and rewilding initiatives (Byers et al., 2006). However, the impact of ecosystem engineers on biodiversity may be more pronounced when considered in a broader context, both temporal and spatial, or integrated with other well-established ecological processes. For example, the temporal outcome of their activities exceeds their life-span because of the legacy effect, and the biogenic structures they have created, such as termite mounds or woodpecker cavities, remain in the landscape and continue to function after their death (Trzcinski et al., 2021; Albertson et al., 2022). In the spatial context, the effects of ecosystem engineering can be expected to extend beyond the area of a species’ immediate activity. Such a change in an ecosystem that has occurred at greater distance from the source of impact is referred to as a spill-over effect, and is exemplified by local protected areas that contribute to increasing biodiversity beyond their boundaries (Brodie et al., 2023). Although many researchers have investigated the impacts of ecosystem engineers on biodiversity in the area where they are active, a species’ spatial impact may extend farther than previously thought when indirect or cascading effects are taken into consideration.

Among the iconic engineering species is the Eurasian beaver *Castor fiber* (henceforth: beaver), appreciated provider of numerous ecosystem services (Law et al., 2019; Brazier et al., 2021). The recently expanding beaver population has been attracting increasing attention as an agent in the restoration of freshwater and riparian ecosystems, which are of high conservation concern throughout the world (Naiman et al., 1993). To a remarkable extent, beavers are capable of transforming the streams they inhabit by creating a heterogeneous mosaic of habitats consisting of a system of newly-formed, mature and abandoned ponds, extending up to several kilometres along a watercourse (Graf et al., 2016; Campbell-Palmer et al., 2021). Such beaver-induced transformations have considerable consequences for channel geomorphology and biogeochemistry, namely, increased retention, improved water quality, reduced erosion and other changes in watercourse properties (Pollock et al., 2018). Modification of water regimes combined with altered vegetation offers rich and diverse habitats for a broad spectrum of organisms (Windels, 2017; Willby et al., 2018). There is a large body of evidence that demonstrates the keystone role of beavers in the ecosystem, as the appearance of this ecological engineer is associated with an increase in the species richness and abundance of water-related taxa (Kemp et al., 2012; Smith and Mather, 2013; Bush et al., 2019; Dalbeck et al., 2020; Washko et al., 2022).

Riparian habitats, situated between terrestrial and aquatic ecosystems, have unique plant and animal assemblages and therefore enhance local species pools (Sabo et al., 2006). Being semi-aquatic mammals, beavers may utilize habitats located up to 100 m from the water’s edge (Stoffyn-Egli and Willison, 2011), where its foraging activities change the composition and structure of vegetation (Johnston and Naiman, 1990; Svanholm Pejstrup et al., 2023). Hence, the presence of beavers and the habitat modifications they bring about can enhance the biodiversity of terrestrial ecosystems adjacent to their ponds as a result of cascading effects (Mourant et al., 2018; Fedyń et al., 2022). Even though boundaries between aquatic and terrestrial ecosystems are obvious, they are readily crossed, e.g. by flows of energy and resources (Marczak et al., 2007). The link between terrestrial and aquatic ecosystems is well exemplified by the key impact of migratory salmon *Onchorhycus* spp. on the functioning of terrestrial ecosystems adjacent to streams (Hocking and Reynolds, 2011; Wagner and Reynolds, 2019) or by the cascading effect of invasive rainbow trout *Oncorhynchus mykiss* resulting in the decline in forest spiders and birds (Baxter et al., 2004; Epanchin et al., 2010). As the activity of beavers directly or indirectly affect many components of aquatic ecosystems (Law et al., 2016; Grudzinski et al., 2022), it is anticipated that diverse biotic and abiotic changes in them may also have consequences for adjoining terrestrial ecosystems. Biodiversity loss is of global concern and affects a great many taxa and habitats (Dirzo et al., 2014), so the extension of “beaver keystones” beyond the aquatic zone will have conservation implications for the restoration of terrestrial ecosystems.

Birds are potential ecological indicators of terrestrial biodiversity. They are a widespread, diversified and mobile group of animals which occupy high levels in the trophic pyramid, and thus respond well to habitat modifications and changes in the composition and abundance of organisms at lower levels of food chains (Gregory and Strien, 2010; Fraixedas et al., 2020). Apart from their indicative value, birds are an important group of organisms as many species are of conservation concern and their populations are locally and globally declining (Gregory et al., 2007; Inger et al., 2015). In the case of beavers, it has been shown that the wetlands they create host more species and individuals of breeding or wintering birds than areas undisturbed by these animals (Grover and Baldassarre, 1995; Aznar and Desrochers, 2008; Orazi et al., 2022; Fedyń et al., 2023). However, the spatial range of the effect of beavers on breeding birds along the wetland-forest gradient remains unknown.

Although there is much evidence pointing to the important ecological role of beaver wetlands, the spatial range of the “beaver effect” on terrestrial biodiversity remains unclear. This study explores the species richness, abundance and assemblage composition of breeding birds at increasing distances from the water’s edge along watercourses modified by beavers and compares these with bird assemblages along unmodified ones. We test the hypothesis that the engineering activity of beavers induces a biodiversity spill-over effect on to adjacent terrestrial habitats. We expect that beaver wetlands are hot-spots of breeding bird richness and abundance, and that the effects of the beaver’s presence and activities are also reflected in the surrounding terrestrial habitats. This would highlight the cross-ecosystem consequences of ecosystem engineering.

## Methods

### Study area

This survey was conducted in Poland (central Europe). This country lies in the temperate zone with a mean annual air temperature of 9.9°C and an average annual precipitation of 645 mm. The average altitude in Poland is 173 m amsl, and the topography is diverse, ranging from mountains (the highest peak is 2499 m amsl) to lowlands. The country is dominated by farmland (60%) and forests (30%), while urbanized areas make up just 5% of the area.

The study sites were situated in lowland, upland and mountain regions in a range of forest habitats that can be inhabited by Eurasian beaver in the temperate climatic zone (Fig. 1). The lowland areas were located on the Northern Podlasie Plain (average altitude 200 m amsl; NE Poland) in the Białowieża Forest and Knyszyn Forest, extensive complexes dominated by deciduous woodland with European hornbeam *Carpinus betulus* and pedunculate oak *Quercus robur* and coniferous stands with Scots pine *Pinus sylvestris*. The uplands (altitudes 200 – 400 m amsl) included the Sandomierz Basin and Kielce Upland, where the land cover is a mosaic of farmland and forests with Scots pine or silver fir *Abies alba*. The mountain areas surveyed were in the Beskidy Mountains in the Carpathians (altitudes 300 – 600 m amsl), where the forests are dominated by European beech *Fagus sylvatica* and silver fir. Throughout the country, riparian habitats along watercourses are naturally dominated by alder *Alnus* spp., willow *Salix* spp. and poplar *Populus* spp. Although the size of water-related habitat patches is largely a consequence of the land configuration, the state of preservation of such habitats depends how the surrounding land is used.

**Fig. 1.**
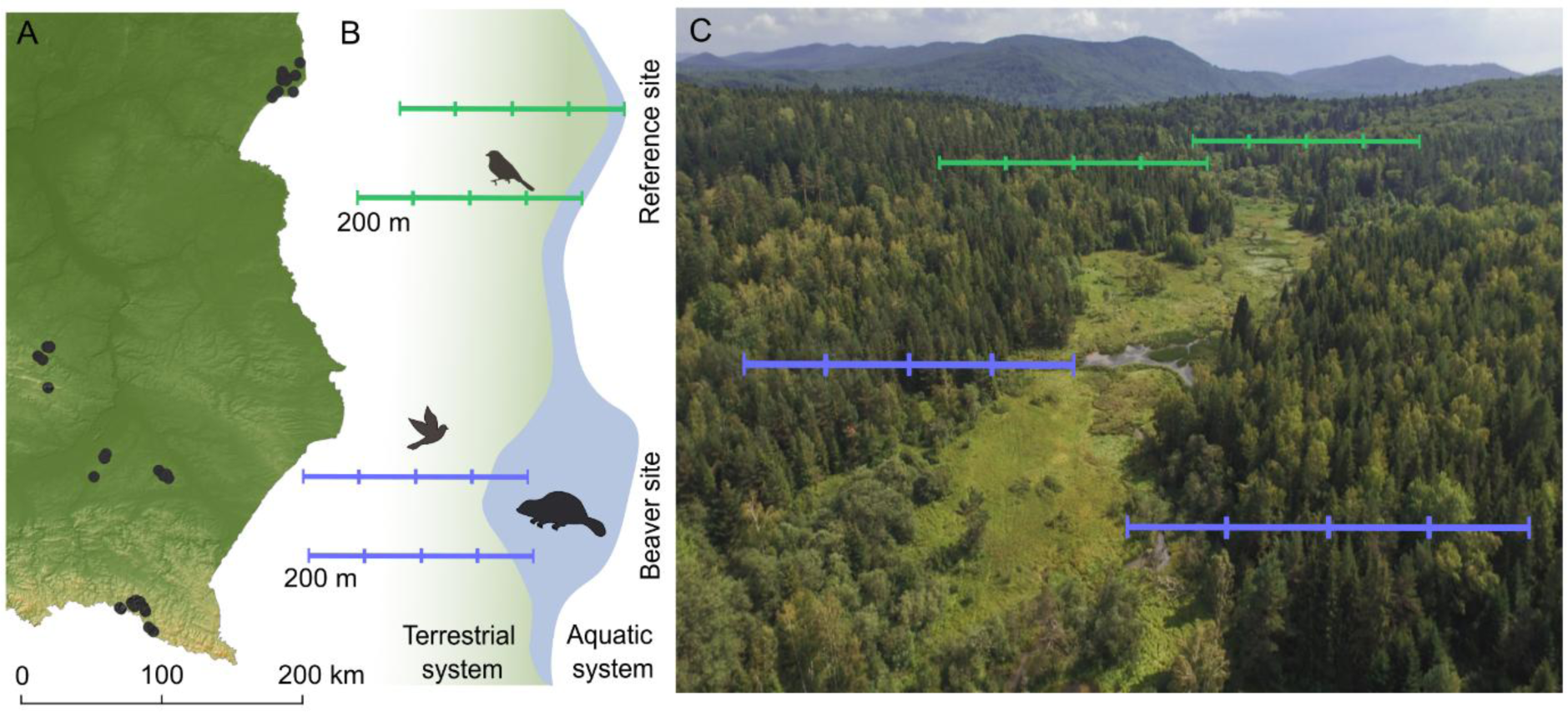
Map of the study area (Poland, central Europe) showing the positions of pairs (N = 40) of Eurasian beaver *Castor fiber* sites and reference sites (A) and the study scheme (B); the photograph (C) shows a typical research site with the transects where birds were counted.

As in many parts of Europe, beaver populations in Poland have been recovering over the last few decades following reintroductions and natural expansion (Żurowski and Kasperczyk, 1988). After World War II, beavers were thought to be extinct in the country, but in 1975 their numbers started to increase, and have done so consistently ever since (Wróbel and Krysztofiak-Kaniewska, 2020). At present, the Polish beaver population of some 120 000 individuals is widely distributed across the country (Halley et al., 2021). Beavers are becoming increasingly common, and they are being recorded in all types of landscapes, from forests to human-modified areas such as urban and agricultural habitats (Ciach et al., 2023).

### Field methods

#### Study site selection

The survey was carried out on 80 study sites, consisting of 40 pairs of Eurasian beaver sites (hereafter – beaver sites) and sites not inhabited by this ecosystem engineer (hereafter – reference sites) (Fig. 1). Thirty-four of the beaver sites were situated in parts of watercourses transformed by these animals into wetlands by the impoundment of water (dam building). On the other 6 sites, beavers had colonized existing wetlands, i.e. natural old riverbeds. The beaver sites surveyed were situated in forest interiors, among bankside woody vegetation along watercourses, or in ecotone habitats on forest margins. The beaver sites ultimately included in the study, which had to be at least 100 m long, were randomly selected from the pre-prepared database of sites occupied by the species in the study area. The mean length of the study sites was 523.48 ± 354.90 SD (range 110 – 1500) m and their mean area was 1.60 ± 2.17 SD ha (range 0.15 – 10.50). Stretches of watercourses without signs of present or past beaver activity were designated as reference sites and these lay upstream or downstream of the beaver sites (Fig. 1). The reference sites were situated in a similar landscape context as their paired beaver sites, i.e. if a beaver site was located within a forest, the reference site was also situated in such a habitat. The reference sites included in the study lay at average distances of 811.86 m ± 845.68 SD (range 200 – 3000) from the beaver sites and were located on watercourses which had an average width of 1.56 m ± 0.82 SD (range 0.5 – 4 m).

We considered beaver wetlands to be parts of watercourses modified by the beavers’ activity: active or abandoned ponds with adjacent grassy and shrubby vegetation that emerged as a result of the animals’ flooding or foraging activity. The total area of a beaver site was measured along the total length of the beaver-modified part of the watercourse, including all the habitats that had come into being as a result of the animals’ activity. We took a single beaver site to be an area with numerous signs of beaver activity (ponds and adjacent areas with modified vegetation). Two areas with signs of beaver activity separated by a distance of more than 500 m were considered to be separate sites.

#### Bird sampling

In order to assess the differences in the bird assemblages between the beaver and reference sites along a gradient of increasing distance from the water’s edge, birds were counted on transects (Fig. 1). Two 200 m long transects were laid out at right angles to the water’s edge on each beaver and reference site. The transects were set up at random, with a distance of 200 m between them on a given beaver / reference site. The transects were divided into four 50 m long sections (hereafter – distance zones). The starting point of the first section was at the water’s edge, and included 25 m of the water area and 25 m of the land area. The observer recorded all birds seen or heard up to 100 m from the transect. Birds flying over the transects were recorded separately. The transects within one pair of study sites (beaver and reference) were sampled on one morning in random order. Each site was visited twice: once in May and again in June 2021 during the early morning hours from dawn to 10:00 hrs. The counts were done on days with favourable weather conditions, i.e. without rain, strong winds (wind speed ≤ 10 km/h) or fog. The field surveys were carried out by four experienced field ornithologists.

### Data handling and analyses

A total of 6,465 birds were recorded during the two field surveys. Prior to the analyses, birds noted >100 m from the transect axis and in flight were excluded from the dataset. Then, the number of individuals of each species in each distance zone (50 m long, 200 m wide) on each transect was determined as the maximum number of individuals of a given species recorded during the two surveys (N = 160). The final dataset included 4,677 bird records. Within each study plot (N = 80), the numbers of individuals of each species were summed from the two transects within the several distance zones (< 50, 50 – 100, 100 – 150 and 150 – 200 m). The occurrence of each species (binomial variable – species present or absent on a given site) was determined for each distance zone on each site for both transects combined. All further analyses were based on the numbers of individuals/occurrence summed from the two transects to the site level in each distance zone. The total number of individuals was calculated as the sum of the individuals of all species detected within the distance zones on each site. The total number of species was calculated as the sum of all species occurring in the distance zones on each site.

ANOVA with Tukey’s post-hoc HSD test for multiple comparisons were performed to determine the differences between the means of the total number of species and individuals on the beaver and reference sites in each distance zone. The frequency of occurrence of a given bird species on the beaver and reference sites in each distance zone were calculated as the fraction of sites where the species was present. Differences between the frequency of occurrence and the total number of individuals of each species were tested using the chi-square test with Yates’ correction and the Mann-Whitney U test, respectively. A minimum probability level of p < 0.05 was considered significant. Results with p < 0.06 were considered as approaching significance level. Owing to the predicted nonlinear relationship between the area of a beaver site and the distance from the water and the total number of species / individuals, generalized additive models (GAMs) were used. The ‘mgcv’ package implemented in R was used to create and visualize GAMs (Wood, 2017).

The numbers of individuals of each species on each study site (N = 80) in every distance zone on the beaver and reference sites were used for the avian community analyses. Prior to the analyses, the number of individuals of each species was standardized to relative abundance in order to moderate the effect of common species for analysis and visualization. Following the procedure suggested by Buttigieg and Ramette (2014), analysis of similarity (ANOSIM) was performed on the basis of the Bray-Curtis dissimilarity matrix to test differences in species composition between the beaver and reference sites in each distance zone. Non-metric multidimensional scaling (NMDS) was employed to generate graphical representations of the bird assemblages (abundances of all species observed) recorded on the beaver and reference sites in all the distance zones. The two-dimensional ordination was based on the Bray-Curtis dissimilarity matrix. The initial two-dimensional ordination generated stress values approaching 0.27, indicating a low ordination fit. In order to reduce stress, a three-dimensional ordination was run, but as this did not sufficiently reduce the stress value (0.24) and rendered plot interpretation less intuitive, it was decided to perform the two-dimensional visualization. Community analyses were performed using the ‘vegan’ package (Oksanen, 2021) in R (R Core Team, 2022).

The IUCN Red List Category and trends for European populations were used (BirdLife International, 2021) to confirm the conservation status of species whose frequency of occurrence or numbers of individuals were affected by beavers. The European Red List of Birds is based on data reported by Member States of the European Union in line with the methodological paper by Röschel et al. (2020). Data standardized with the recommendations of the IUCN Red List Categories and Criteria are used as basic indicators of biodiversity. The assessment of such a regional extinction risk identifies priorities and helps to inform decision makers in conservation policy (Rodrigues et al., 2006).

## Results

A total of 106 bird species were recorded. The overall number of species detected on the beaver sites was 28.6% higher than on the reference ones, 99 and 77, respectively. Likewise, the mean total number of species detected on the beaver sites was higher than on the reference sites (23.3 ± 5.7 SD *vs* 18.6 ± 3.4 SD; W = 1214, p < 0.000). The mean total number of individuals on the beaver sites was 67.4 ± 25.8 SD, while on the reference sites it was 49.5 ± 17.2 SD (W = 1203, p < 0.000).

The total number of species and the total number of individuals differed between the beaver and reference sites and between the distance zones within the beaver or reference sites (Fig. 2). On both the beaver and reference sites, the total number of species and the total number of individuals were the highest near the water and decreased with increasing distance from the water’s edge (Fig. 2). There were higher total numbers of species (p = 0.000) and of individuals (p = 0.000) on the beaver sites than on the reference sites in the < 50 m distance zone (Fig. 2). In the 50 – 100 m distance zone, the total numbers of species (p = 0.046) and of individuals (p = 0.036) on the beaver sites were higher than on the reference sites (Fig. 2). In the 100 – 150 distance zone there were no differences between the beaver and reference sites as regards the totals number of species (p = 0.776) or of individuals (p = 0.669; Fig. 2). Likewise, there were no differences in the 150 – 200 m distance zone between the beaver and reference sites in the total numbers of species and of individuals (p = 0.814 and p = 0.974, respectively; Fig. 2).

**Fig. 2.**
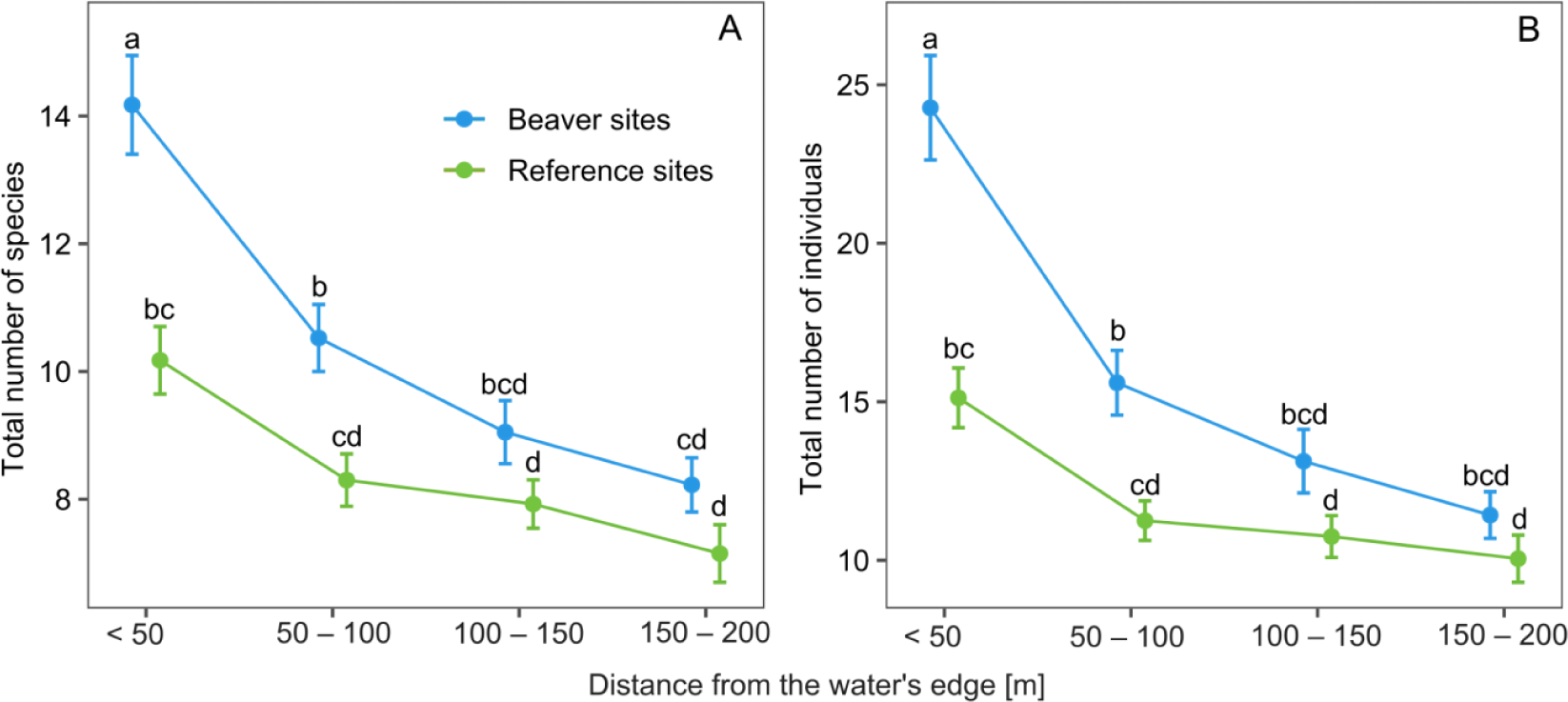
Mean (dots) and SE (whiskers) of the total number of species (A) and the total number of individuals (B) recorded on transects divided into distance zones (< 50 m, 50 – 100 m, 100 – 150 m and 150 – 200 m) from the water’s edge on Eurasian beaver *Castor fiber* sites (N = 40) and paired reference sites (N = 40) during the breeding season in Poland (central Europe). Differences were tested with ANOVA and Tukey’s post-hoc test for multiple comparisons. The same lower-case letters indicate the absence of a statistical difference between the means at p < 0.05.

26.9% of the recorded species occurred exclusively on the beaver sites (Table S1). Water-related species like garganey *Spatula querquedula*, grey heron *Ardea cinerea*, common goldeneye *Bucephala clangula*, black stork *Ciconia nigra*, marsh harrier *Circus aeruginosus* and water rail *Rallus aquaticus* were found only on the beaver sites (Table S1). Moreover, reed-and shrub-dwelling birds like sedge warbler *Acrocephalus schoenobaenus*, marsh warbler *Acrocephalus palustris*, river warbler *Locustella fluviatilis*, Savi’s warbler *Locustella luscinioides*, whitethroat *Curruca communis* and thrush nightingale *Luscinia luscinia* were likewise recorded only on the beaver sites (Table S1). Birds foraging over water, such as sand martin *Riparia riparia*, barn swallow *Hirundo rustica* and common swift *Apus apus*, were also found exclusively at the beaver sites. Some cavity nesting species, such as pygmy owl *Glaucidium passerinum*, grey-headed woodpecker *Picus canus*, tawny owl *Strix aluco*, hoopoe *Upupa epops* and short-toed treecreeper *Certhia brachydactyla*, occurred exclusively on the beaver sites (Table S1). The forest-dwelling species that were found more frequently on the beaver sites included tree pipit *Anthus trivialis*, collared flycatcher *Ficedula albicollis*, white-backed woodpecker *Dendrocopos leucotos*, jay *Garrulus glandarius* and blackcap *Sylvia atricapilla* (Table S1).

The composition of the breeding bird assemblage differed between the beaver and reference sites in the < 50 m and 50 – 100 m distance zones (Fig. 3). At greater distances from the water’s edge (the 100 – 150 m and 150 – 200 m distance zones) there were no differences in the composition of the bird assemblage (Fig. 3). The number of species detected exclusively on the beaver sites was higher than on the reference sites in the < 50 m and 50 – 100 m distance zones (Fig. 3). In the < 50 m zone, nine species occurred more frequently on the beaver sites than on the reference sites, five of which were species with decreasing population trends in Europe (Table 1). There were 13 species which were more abundant in the < 50 distance zone, including seven species with decreasing population trends in Europe, and the difference in the abundance of two other species approached significance (Table 2). Four species occurred more frequently on the beaver sites than on the reference sites in the 50 – 100 m distance zone, including two with decreasing population trends (Table 1). Seven species were more abundant on the beaver sites than on the reference sites in 50 – 100 m distance zone, including five with decreasing population trends in Europe (Table 2). There were no species that occurred more frequently on the beaver sites or reference sites in the 100 – 150 m zone, but the frequency of occurrence of one species did approach significance in the 150 – 200 m distance zone (Table 1). In the 100 – 150 m distance zone, three species were more abundant on the beaver sites than on the reference sites (Table 2). In the 150 – 200 m distance zone, three species were more abundant on the beaver sites (including one species with decreasing population trends in Europe) and two were more abundant on the reference sites (Table 2).

**Fig. 3.**
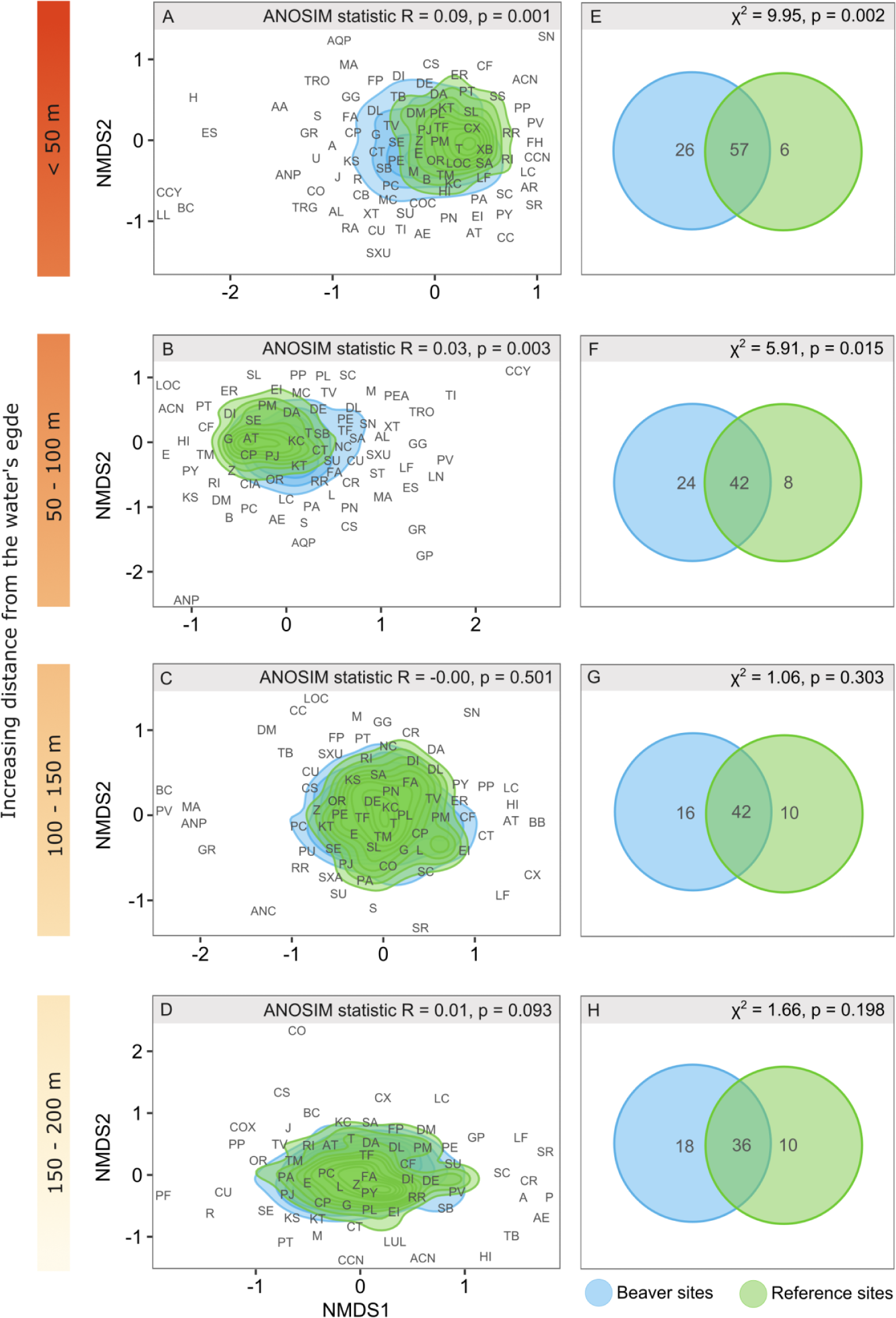
Similarities of the bird assemblage composition on Eurasian beaver *Castor fiber* sites (N = 40) and paired reference sites (N = 40) in distance zones (A – < 50, B – 50 – 100, C – 100 – 150 and D – 150 – 200 m) from the water’s edge and the number of species unique to either the beaver or reference sites or common to both in the distance zones (E – < 50, F – 50 – 100, G – 100 – 150 and H – 150 – 200 m) during the breeding season in Poland (central Europe). This graphic visualization of the bird assemblage was generated using non-metric multidimensional scaling (NMDS), and the differences in species composition between the beaver and reference sites in each distance zone were analysed on the basis of the Bray-Curtis dissimilarity matrix using the analysis of similarity (ANOSIM). The differences in the number of unique species between the beaver and reference sites were tested with the chi-square test. The letters indicating the species codes are explained in Table S1.

**Table 1.**
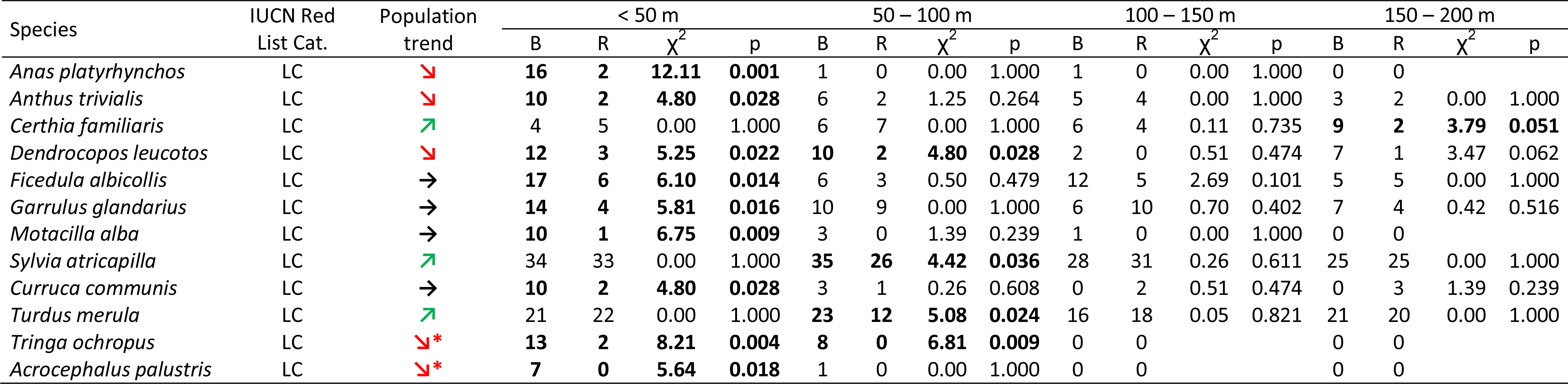
List of bird species whose occurrence (number of sites where the species was recorded) differs between the Eurasian beaver *Castor fiber* sites (B) and reference sites (R) in Poland (central Europe) in four distance zones (< 50 m, 50 – 100 m, 100 – 150 m and 150 – 200 m) from the water’s edge (for the full list of species, see Table S1). IUCN Red List Category for European populations and population trend **↘**-decreasing, **↗**-increasing, **→**-stable, *-data for EU28 region if European data unavailable (BirdLife International, 2021). Differences in frequency of occurrence between beaver and reference sites were chi-square tested. Differences that are significant (p < 0.05) or approaching significance (p < 0.06) are shown in bold.

| Species | IUCN Red List Cat. | Population trend | < 50 m |  |  |  | 50 – 100 m |  |  |  | 100 – 150 m |  |  |  | 150 – 200 m |  |  |  |
| --- | --- | --- | --- | --- | --- | --- | --- | --- | --- | --- | --- | --- | --- | --- | --- | --- | --- | --- |
| | | | B | R | $\chi^2$ | p | B | R | $\chi^2$ | p | B | R | $\chi^2$ | p | B | R | $\chi^2$ | p |
| *Anas platyrhynchos* | LC |  | **16** | **2** | **12.11** | **0.001** | 1 | 0 | 0.00 | 1.000 | 1 | 0 | 0.00 | 1.000 | 0 | 0 |  |  |
| *Anthus trivialis* | LC |  | **10** | **2** | **4.80** | **0.028** | 6 | 2 | 1.25 | 0.264 | 5 | 4 | 0.00 | 1.000 | 3 | 2 | 0.00 | 1.000 |
| *Certhia familiaris* | LC |  | 4 | 5 | 0.00 | 1.000 | 6 | 7 | 0.00 | 1.000 | 6 | 4 | 0.11 | 0.735 | **9** | **2** | **3.79** | **0.051** |
| *Dendrocopos leucotos* | LC |  | **12** | **3** | **5.25** | **0.022** | **10** | **2** | **4.80** | **0.028** | 2 | 0 | 0.51 | 0.474 | 7 | 1 | 3.47 | 0.062 |
| *Ficedula albicollis* | LC |  | **17** | **6** | **6.10** | **0.014** | 6 | 3 | 0.50 | 0.479 | 12 | 5 | 2.69 | 0.101 | 5 | 5 | 0.00 | 1.000 |
| *Garrulus glandarius* | LC |  | **14** | **4** | **5.81** | **0.016** | 10 | 9 | 0.00 | 1.000 | 6 | 10 | 0.70 | 0.402 | 7 | 4 | 0.42 | 0.516 |
| *Motacilla alba* | LC |  | **10** | **1** | **6.75** | **0.009** | 3 | 0 | 1.39 | 0.239 | 1 | 0 | 0.00 | 1.000 | 0 | 0 |  |  |
| *Sylvia atricapilla* | LC |  | 34 | 33 | 0.00 | 1.000 | **35** | **26** | **4.42** | **0.036** | 28 | 31 | 0.26 | 0.611 | 25 | 25 | 0.00 | 1.000 |
| *Curruca communis* | LC |  | **10** | **2** | **4.80** | **0.028** | 3 | 1 | 0.26 | 0.608 | 0 | 2 | 0.51 | 0.474 | 0 | 3 | 1.39 | 0.239 |
| *Turdus merula* | LC |  | 21 | 22 | 0.00 | 1.000 | **23** | **12** | **5.08** | **0.024** | 16 | 18 | 0.05 | 0.821 | 21 | 20 | 0.00 | 1.000 |
| *Tringa ochropus* | LC |  | **13** | **2** | **8.21** | **0.004** | **8** | **0** | **6.81** | **0.009** | 0 | 0 |  |  | 0 | 0 |  |  |
| *Acrocephalus palustris* | LC |  | **7** | **0** | **5.64** | **0.018** | 1 | 0 | 0.00 | 1.000 | 0 | 0 |  |  | 0 | 0 |  |  |

**Table 2.**
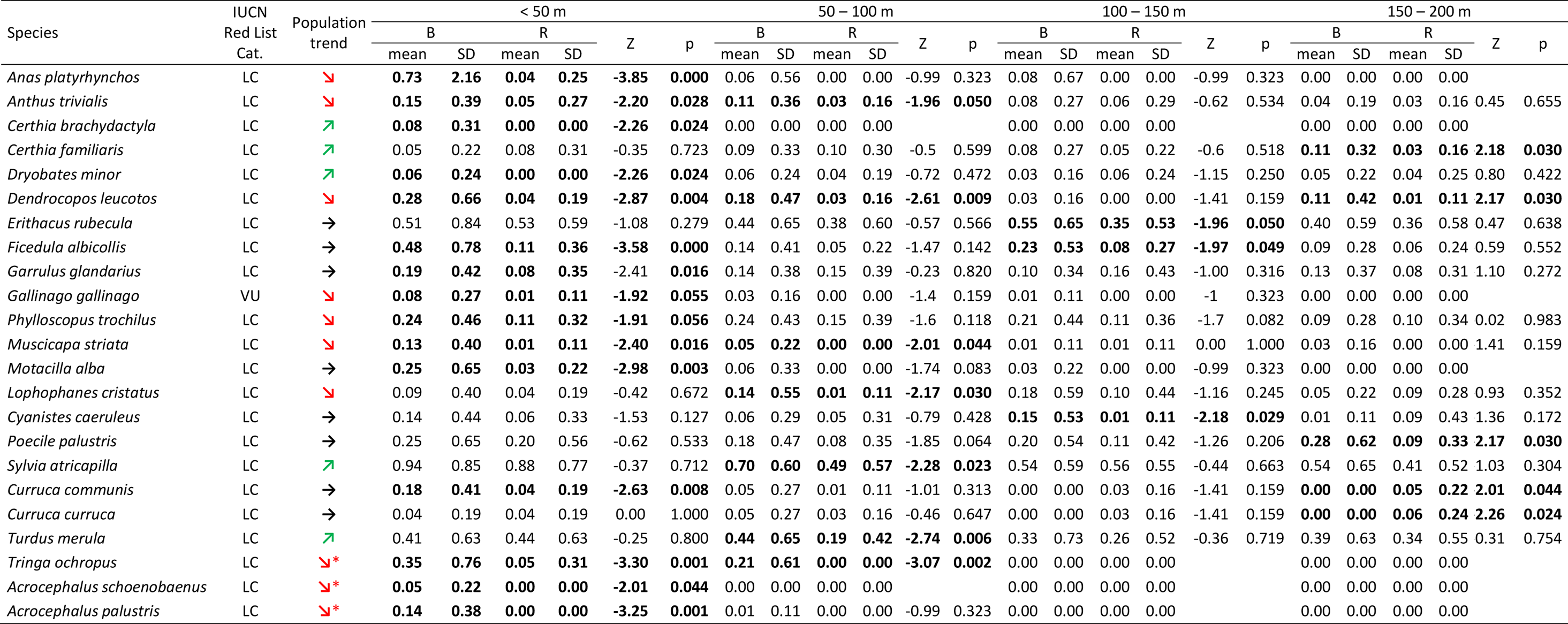
List of bird species whose mean number of individuals per 100 m of transect differs between the Eurasian beaver *Castor fiber* sites (B) and reference sites (R) in Poland (central Europe) in four distance zones (< 50 m, 50 – 100 m, 100 – 150 m and 150 – 200 m) from the water’s edge (for the full list of species, see Table S2). IUCN Red List Category for European populations and population trend **↘**-decreasing, **↗**-increasing, **→**-stable, *-data for EU28 region if European data unavailable (BirdLife International, 2021). Differences in the numbers of individuals between beaver and reference sites were tested with the Mann Whitney U test. Difference that are significant (p < 0.05) or approaching significance (p < 0.06) are shown in bold.

The total number of species and the total number of individuals on the beaver sites were correlated positively with the area of beaver wetland but negatively with increasing distance from the water’s edge (Table 3, Fig. 4). The area of wetland and the distance from the water both explained 42.1% of the deviance in the total number of species and 41.1% in the total number of individuals (Table 3). Although the numbers of species and individuals decreased with increasing distance from the water’s edge in wetlands of all sizes, the largest wetlands in the 150 – 200 m distance zone from the water had as many species and individuals as the smallest wetlands in the < 50 m distance zone (Fig. 4).

**Fig. 4.**
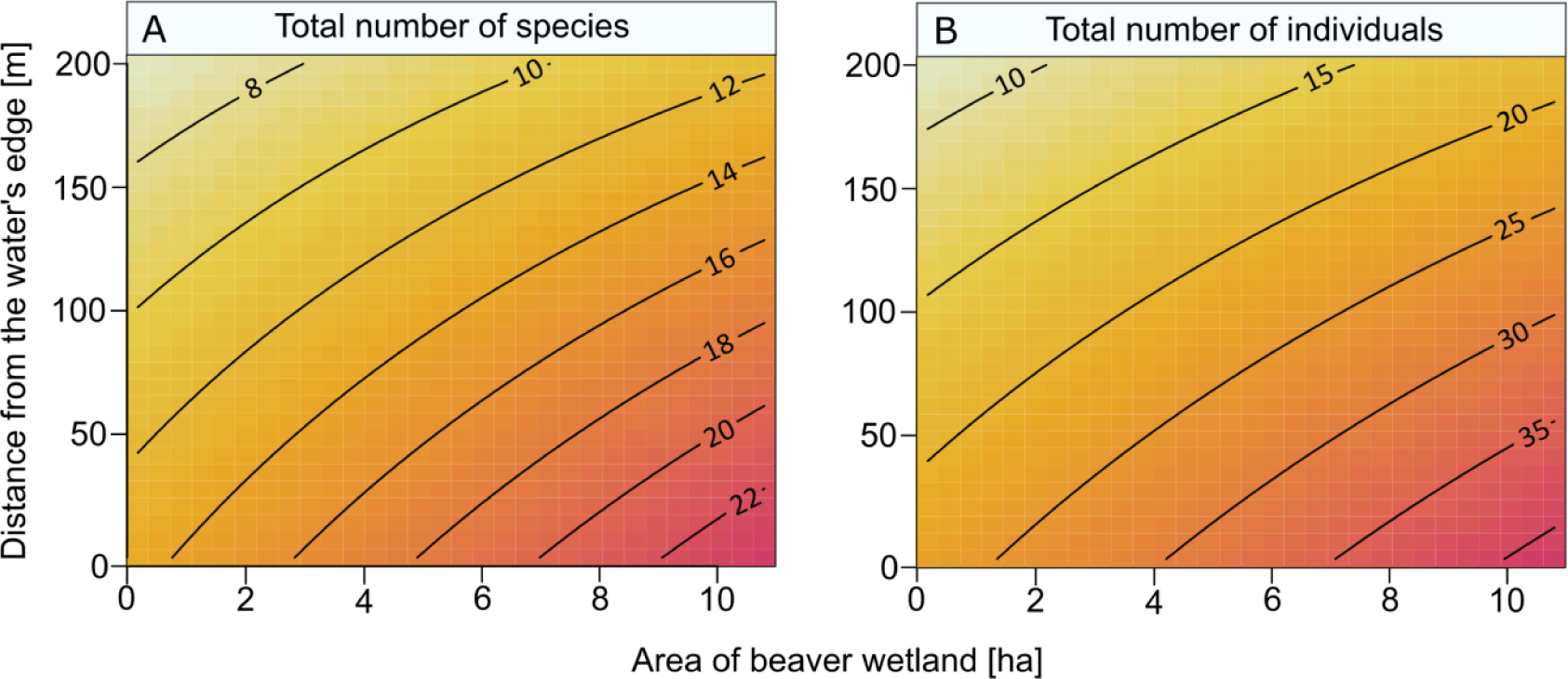
The total number of species (A) and the total number of individual birds (B) on Eurasian beaver *Castor fiber* sites (N = 40) during the breeding season in Poland (central Europe) in relation to the area of beaver wetlands and the distance from the water’s edge, as predicted by the generalized additive models shown in Table 3.

**Table 3.** Summary of generalized additive models explaining the total number of species and the total number of individuals per 100 m of transect on the Eurasian beaver *Castor fiber* sites (N = 40) during the breeding season in Poland (central Europe) in relation to the area of beaver wetland and distance from the water’s edge.

| Response variable | Explanatory variables | Estimate | SE | t | p | Deviance explained (%) |
| --- | --- | --- | --- | --- | --- | --- |
| Total number of species | Intercept | 13.93 | 0.66 | 21.01 | 0.000 | 42.1 |
|  | Area | 0.66 | 0.10 | 6.57 | 0.000 |  |
|  | Distance | -0.04 | 0.00 | -8.42 | 0.000 |  |
|  | Intercept | 9.10 | 0.39 | 22.77 | 0.000 | 15.9 |
|  | Area | 0.66 | 0.12 | 5.47 | 0.000 |  |
|  | Intercept | 15.33 | 0.71 | 21.66 | 0.000 | 26.2 |
|  | Distance | -0.04 | 0.01 | -7.48 | 0.000 |  |
| Total number of individuals | Intercept | 23.72 | 1.38 | 17.17 | 0.000 | 41.1 |
|  | Area | 1.25 | 0.21 | 5.98 | 0.000 |  |
|  | Distance | -0.08 | 0.01 | -8.58 | 0.000 |  |
|  | Intercept | 13.47 | 0.84 | 16.08 | 0.000 | 13.4 |
|  | Area | 1.25 | 0.25 | 4.95 | 0.000 |  |
|  | Intercept | 26.36 | 1.45 | 18.23 | 0.000 | 27.6 |
|  | Distance | -0.08 | 0.01 | -7.77 | 0.000 |  |

## Discussion

The results of our study show that beaver wetlands constitute hot-spots of breeding bird species richness and abundance that spill over on to adjacent terrestrial areas. We found that beaver wetlands host higher numbers of species and individual breeding birds than those parts of watercourses unmodified by this ecosystem engineer. Moreover, a significant proportion of the recorded species occurred exclusively at beaver sites, which highlights the importance for local biodiversity of habitats that have come into being as a result of beaver activity. Our findings are in line with previous studies that reported richer and more abundant assemblages of birds on waters modified by beavers than in unoccupied ones (Grover and Baldassarre, 1995; Brown et al., 1996; Aznar and Desrochers, 2008; Orazi et al., 2022). We also found a greater bird species richness and an abundant spill-over on to terrestrial habitats adjacent to beaver wetlands, in contrast to terrestrial habitats located near unmodified watercourses, where there were no such increases in species richness or abundance. The pattern according to which beavers enhance local biodiversity was previously observed in several taxonomic groups, such as plants, birds, bats and small vertebrates (Orazi et al., 2022). Our study builds on this finding and explains how bird richness and abundance change along a gradient from a pond to a forest interior and, thus, address the question of how far the effects of this ecosystem engineer extend across the landscape. We found that breeding bird assemblages were more diverse and abundant up to 100 m from the water than assemblages occupying land adjacent to watercourses unmodified by beavers. As birds are appropriate ecological indicators (Fraixedas et al., 2020), our result suggests that the effects of ecosystem engineering lead to a spill-over of biodiversity on to terrestrial areas adjacent to beaver wetlands.

### Changes in bird assemblages induced by ecosystem engineers

Ecosystem engineering can affect communities of organisms by modifying the environment and thus also the trophic and non-trophic interactions between species (Sanders et al. 2014). Beavers transform running water into stagnant ponds of different ages and plant successional stages (Kivinen et al., 2020). In consequence, beaver sites represent mosaics of heterogeneous habitat patches (Wright and Jones, 2004), offering niches for a greater number of species than streams unmodified by beavers and their adjacent terrestrial habitats (Willby et al., 2018; Bush et al., 2019; Orazi et al., 2022). Although we did not investigate the habitat attributes that specifically affect a bird community, riparian habitats are acknowledged as areas that offer favourable conditions for wildlife owing to the access to water and the beneficial microclimate (Gregory et al., 1991; Bennett et al., 2014). A riparian zone, such as the interface between aquatic and terrestrial ecosystems, potentially provides diverse resources like food, shelter, water and migration corridors (Naiman et al., 1993; Naiman and Decamps, 1997), which can encourage the presence of birds in this habitat. As sunlight is an important driver of freshwater productivity, beaver sites are more productive than streams shaded by tree canopies and thus have richer food resources for birds (Death and Zimmermann, 2005; Hood and Larson, 2014; Washko et al., 2022). Opening the tree canopy changes the temperature, solar radiation, humidity and wind speed along gradients from the water’s edge into the stand interior (Chen et al., 1995; Muscolo et al., 2014); this aspect, in turn, mediates the species composition (Fraver, 1994). In consequence, bird communities in forest canopy gaps of different origin are more diverse and abundant (Fuller, 2000; Moorman et al., 2012; Lewandowski et al., 2021).

This study shows that breeding bird assemblages that have developed under the impact of beavers are more diverse, abundant and differ in species composition. Although the majority of bird species used both the beaver and reference sites, 27% of the recorded species occurred exclusively on the beaver sites, indicating that beavers create unique habitats for birds in temperate zone forests. Beaver sites were found to host both terrestrial and aquatic birds, so these areas clearly offer habitats for species with diverse ecological requirements. Previous studies reported that beaver wetlands support the occurrence of habitat specialists such as waterbirds, which are less likely to occur in landscapes lacking beaver-modified habitats (Nummi and Holopainen, 2014). It was also found that beavers encouraged the presence of forest habitat specialists like white-backed woodpecker, collared flycatcher, pygmy owl, treecreeper and hoopoe, i.e. species dependent on cavities and, in some cases, crucially dependent on coarse woody debris. Riparian habitats provide abundant cavities excavated by primary cavity nesters (Przepióra and Ciach, 2022). Suitable habitats for cavity nesters contain large amounts of coarse woody debris (Lõhmus, 2010), which beavers can provide (Thompson et al., 2016). It has been already shown that beaver wetlands are favourable breeding sites for rare birds associated with dead wood, such as three-toed woodpecker *Picoides tridactylus* (Pietrasz et al., 2019). Moreover, beaver activities give rise to terrestrial habitats with dense successional vegetation (Barnes and Dibble, 1988), and our survey showed that birds associated with shrubby vegetation were indeed found only in the beaver wetlands. Our results demonstrate that beavers affect bird assemblages in a much more complex way than previously believed and suggest that the effects of ecosystem engineers should be considered at broader spatial scales in order to assess the full range of their environmental impact.

### Spatial range and consequences of the beaver-induced spill-over effect

We found that the species richness and abundance of birds decreased with increasing distance from the water to the forest interior. However, the species richness and abundance on the beaver sites were higher not only in the immediate vicinity of the water, but also as far as 100 m away from it. Moreover, the species composition of the bird assemblage was clearly different on the beaver sites, not only in the wetland area directly modified by the beavers but also as far as 100 m from the water’s edge. Such a pattern suggests a possible spill-over effect, as a result of which the consequences of the beavers’ activities are felt beyond their immediate area of occurrence (Gray et al., 2016). Being central place foragers, beavers focus their movements and foraging in close proximity to water (Fryxell and Doucet, 1991). Their activities create a habitat gradient, transitioning from open wetlands, through stands affected by flooding and selective foraging, to undisturbed forest beyond the range of immediate impact. In this case, beaver wetlands, supporting diverse and abundant assemblages of different organisms, can provide resources such as food for the bird populations inhabiting the surrounding areas. Birds in ecosystems adjacent to wetlands can benefit, for example, from insects emerging from water, which enrich food webs in terrestrial habitats owing to their relatively good dispersal abilities, i.e. up to 100 m from the water through the forest (Carlson et al., 2016). The beavers’ creation of a diversity of habitats, both within wetlands and in the surrounding areas, enhances the richness and abundance of the nearby bird populations.

Examples are known of spill-over as cross-habitat movements of organisms, e.g. woodland birds or pollinators that move to adjacent open landscapes, which increase diversity and enhance the flow of beneficial ecosystem services such as pest control or pollination (Boesing et al., 2018; Boesing et al., 2022). Moreover, it is likely that the spill-over of biodiversity into areas adjoining beaver wetlands will lead to a parallel spill-over of ecosystem services. Those provided by beavers include a wide range of water-related benefits, such as water retention and purification or carbon sequestration (Thompson et al., 2021). Therefore, the expansion within the landscape of areas potentially performing wetland-related functions will make an invaluable contribution to mitigating climate change.

### Importance of beaver wetland size

We found a positive correlation between the area of a beaver site and the number and abundance of bird species. According to meta-population theory and species-area relationships, a larger area of habitat patches respectively supports larger populations and a greater species richness (Hanski and Ovaskainen, 2000). Increasing areas of wetland may provide more habitats that meet the requirements of open-habitat specialists. Moreover, a larger beaver wetland offers a greater area that is less affected by edge effects (Laurance and Yensen, 1991), which further enhances the quality of the open and wetland habitats thus created.

The area of beaver wetlands increases with the period of occupation of the territory because new impoundments are successively constructed (Kivinen et al., 2020). Therefore, larger beaver sites are probably more heterogeneous than smaller ones, since they consist of ponds of different ages, which differ in the volume of water stored, the successional phase, vegetation composition or deadwood abundance. There is a general relationship between bird diversity and habitat heterogeneity (Bar-Massada and Wood, 2014; Stirnemann et al., 2015), as a diversity of habitats attracts a wide range of bird species with different habitat preferences (Batáry and Báldi, 2004).

Our work adds to the general understanding of the species–area relationship (MacArthur and Wilson, 1967). The shape of the curve describing the species–area relationship depends on covariates (Williams et al., 2009; Tjørve et al., 2021): this is potentially ecosystem-engineer-dependent and may change along with the presence of ecosystem engineers within the pool of species hosted by a given habitat patch. Based on our study, we expect that the species richness in a habitat patch of a given size with an ecosystem engineer will be greater than in such a patch without an ecosystem engineer because the engineering activity extends beyond the patch limits. However, this hypothesis needs further investigation and empirical testing.

### Implication for the conservation of beaver sites as biodiversity hot spots

Riparian habitats are globally important for biodiversity (Sabo et al., 2006; Capon et al., 2013) and host unique bird species at the landscape scale (Bennett et al., 2014). Our study provides important evidence that beavers can play an important role in bird conservation in both wetland and terrestrial ecosystems. We have found that beaver sites are important breeding habitats for species of conservation concern, including species listed in Annex I of the Birds Directive such as white-backed woodpecker and collared flycatcher. Therefore, it is very likely that the beaver’s growing prevalence will have a positive impact on the populations of habitat specialists through the increased availability of habitats created as a result of this ecosystem engineer’s activities. Our study shows that not only habitat specialists or rare species benefit from the beaver’s presence, but also that common ones are more abundant on beaver sites. This is particularly important, given that most of the declines in the European avifauna are attributed to negative population trends of common species (Inger et al., 2015). Therefore, increasing the total number of birds at beaver sites may mitigate overall negative trends of bird abundance (Gregory et al., 2023). Moreover, larger populations of common species have significant ecological consequences, since abundant species make a major contribution to ecosystem functioning (Winfree et al., 2015). Ecosystem engineering performed by various species can also support ecosystem functions by increasing overall biodiversity (Losapio et al., 2023). As birds are indicators of biodiversity (Fraixedas et al., 2020), beavers are also expected to have a positive impact on other groups of terrestrial organisms, additionally underscoring the ecosystem-level effect of engineering.

Currently, beavers are protected in most of their range, and in some places have been reintroduced as a nature-based solution for the restoration of ecosystem services (Brazier et al., 2021; Thompson et al., 2021). The growing beaver population is creating a network of biodiversity hot-spots and may have positive conservation implications at a large spatial scale, overlapping with the species’ range. There is a high potential for engineering effects on ecosystems in areas with viable beaver populations responsible for habitat transformation (Briones, 2024). Moreover, the engineering activities of beavers enhance naturalness and resilience towards disturbances in river valley ecosystems (Fairfax and Whittle, 2020). Our findings show that in order to benefit from the entire spectrum of services that beavers provide it is essential to preserve not only the habitats directly transformed by this ecosystem engineer, but also the areas adjoining them. Recommendations for creating buffer zones around beaver sites have been published in North America, where the beaver has been identified as an umbrella species for riparian obligate animals (Stoffyn-Egli and Willison, 2011), and our results endorse this approach. The beaver’s features, such as its relative frequency and large population size, its well-known biology, its occupancy of a fairly narrow range of habitats, and its sensitivity to human threats (e.g. transformation of watercourses) and ease of monitoring, fully support this ecosystem engineer as an excellent umbrella species, the protection of which benefits large numbers of naturally co-occurring species (Roberge and Angelstam, 2004, Seddon and Leech 2008). In light of widespread declines in biodiversity, beavers have a great potential for large-scale restoration, not only of natural or semi-natural riparian habitats but also of human-modified habitats like agricultural land or green spaces in the urban landscape (Law et al., 2017; Bailey et al., 2019; Ciach et al., 2023).

## Conclusion

The potential of ecosystem engineers is increasingly being acknowledged as supporting biodiversity and ecosystem functioning from local to global scales. We provide evidence for a biodiversity spill-over effect induced by one such ecosystem engineer – the Eurasian beaver – and its conservation value from the cross-system perspective. This study shows that beavers directly support not only freshwater but also terrestrial ecosystems, since the positive effects of the beaver’s activities spread on to terrain adjacent to areas it has already modified. The heterogeneous habitats thus created, from open waters to woodlands and the transitional zone in between, support birds occupying various niches. Beavers not only facilitate the occurrence and abundance of bird species of conservation concern, i.e. rare or declining ones, but also enhance the abundance of common ones, which make major contributions to ecosystem functioning. Our findings emphasize the high priority that should be given to the preservation of networks of beaver sites in the landscape because of the essential services and functions these animals provide. Moreover, our study indicates applied outcomes like the positive relationship between the increasing area of beaver wetlands and biodiversity. Integrating knowledge about the effects of ecosystem engineers with conservation policy can inform future efforts in nature conservation and restoration. This research constitutes a guideline for the European Union’s 2030 Biodiversity Strategy, which recommends the identification and exclusion of the most valuable forests from management. We therefore suggest areas that intended for conservation should include not only habitats directly transformed by beavers, but also woodlands located beyond the sites occupied by this ecosystem engineer.

## Acknowledgements

We wish to express our gratitude to Marek Skruch for his help with the fieldwork. This study was financially supported by the National Science Centre, Poland (grant No. 2021/41/N/NZ8/02423). Michał Ciach was supported by grant No. 2021/41/B/NZ8/03456 from the National Science Centre, Poland.

## Ethical statement

The study was performed in accordance with Polish law.

## Supplementary materials

**Table S1.**
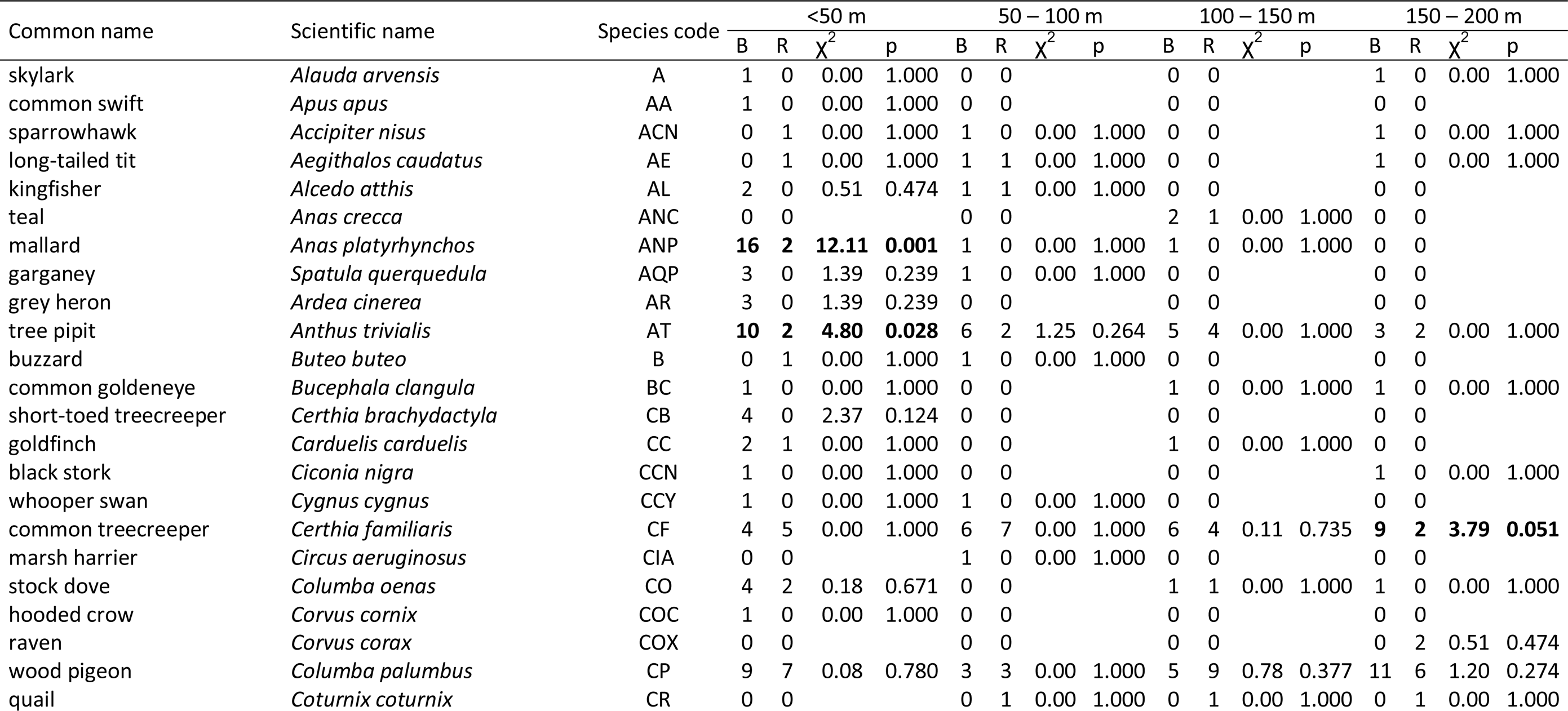
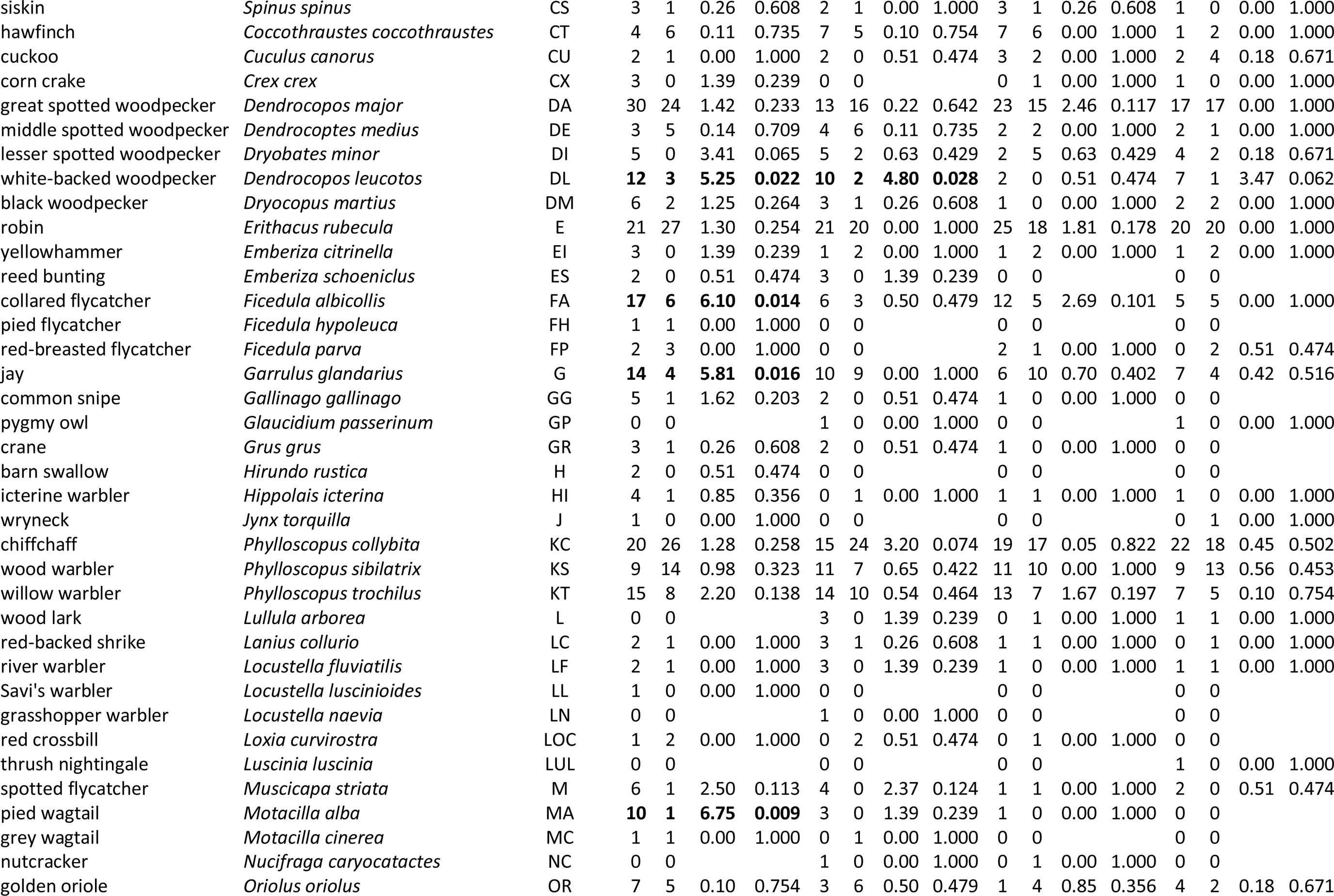
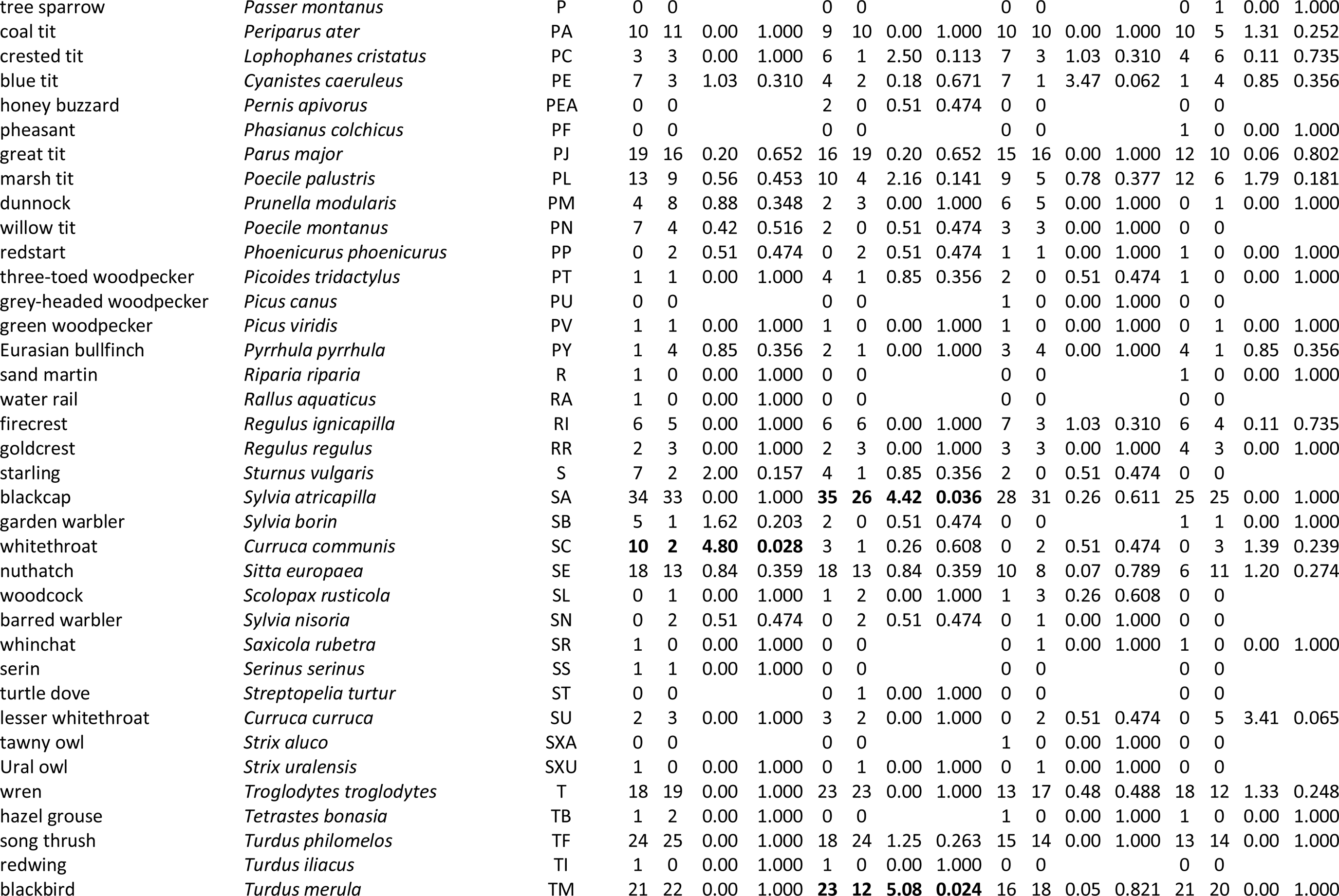
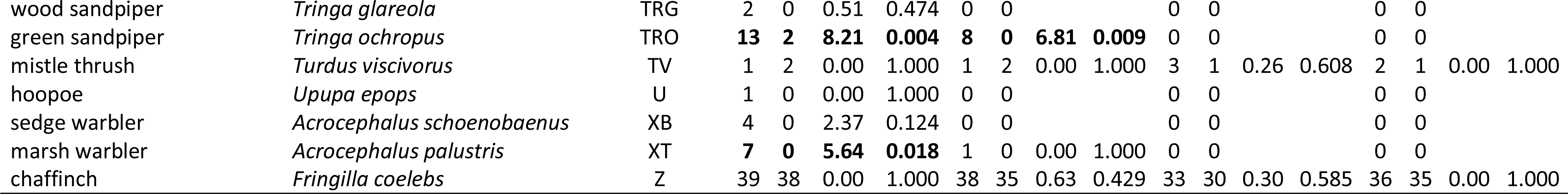
The occurrence of bird species (the number of sites where the species were recorded) on the Eurasian beaver *Castor fiber* sites (B) and reference sites (R) in Poland (central Europe) in four distance zones (< 50 m, 50 – 100 m, 100 – 150 m and 150 – 200 m) from the water’s edge. Differences in the frequency of occurrence between beaver and reference sites were chi-square tested. Differences that are significant (p < 0.05) or approaching significance (p < 0.06) are shown in bold. The species whose frequency differs significantly between the sites are listed in Table 1.

**Table S2.**
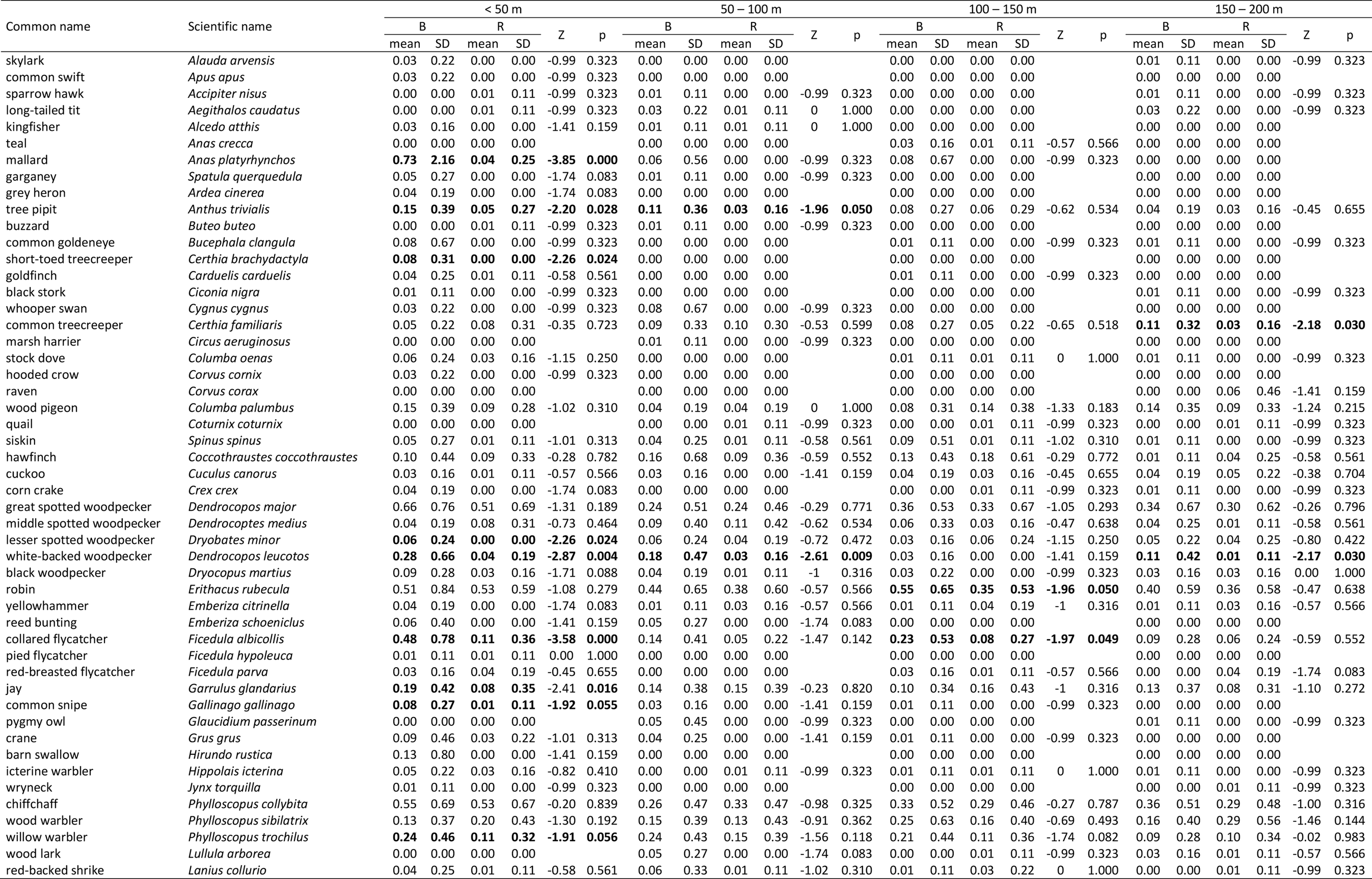
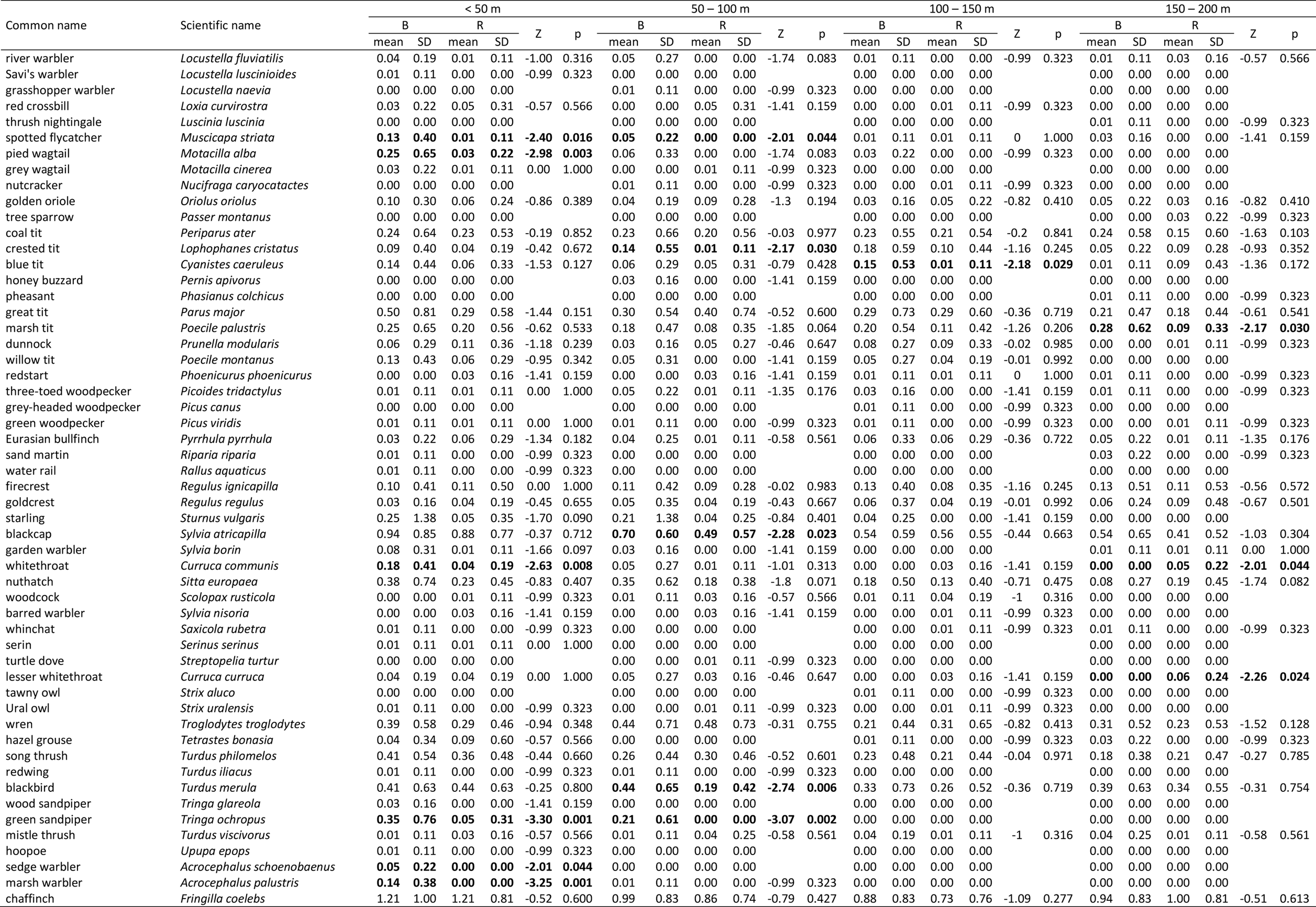
The mean number of individual birds per 100 m of transect recorded on Eurasian beaver *Castor fiber* sites (B) and reference sites (R) in Poland (central Europe) in four distance zones (< 50 m, 50 – 100 m, 100 – 150 m and 150 – 200 m) from the water’s edge. Differences in the number of individuals between the beaver and reference sites were tested with the Mann Whitney U test. Differences that are significant (p < 0.05) or approaching significance (p < 0.06) are shown in bold. The species whose abundance differs significantly between the sites are listed in Table 2.

